# Sequence-function Relationships in Phage-encoded Bacterial Cell Wall Lytic Enzymes and their Implications for Phage-derived Products Design

**DOI:** 10.1101/2021.02.23.432618

**Authors:** Roberto Vázquez, Ernesto García, Pedro García

## Abstract

Phage (endo)lysins are thought to be a viable alternative to usual antibiotic chemotherapy to fight resistant bacterial infections. However, a landscape view of lysins’ structure and properties regarding their function, with an applied focus, is somewhat lacking. Current literature suggests that specific features typical of lysins from phages infecting Gram-negative bacteria (G−) (higher net charge, amphipathic helices) are responsible for an improved interaction with G− envelope. Such antimicrobial peptide (AMP)-like elements are also of interest for antimicrobial molecules design. Thus, this study aims to provide an updated view on the primary structural landscape of phage lysins to clarify the evolutionary importance of several sequence-predicted properties, particularly for the interaction with the G− surface. A database of 2,182 lysin sequences was compiled, containing relevant information such as domain architectures, data on the phages’ host bacteria and sequence-predicted physicochemical properties. Based on such classifiers, an investigation on the differential appearance of certain features was conducted. Such analyses revealed different lysin architectural variants that are preferably found in phages infecting certain bacterial hosts. Particularly, some physicochemical properties (higher net charge, hydrophobicity, hydrophobic moment and aliphatic index) were associated to G− phage lysins, appearing specifically at their C-terminal end. Evidences on the remarkable genetic specialization of lysins regarding the features of the bacterial hosts have been provided, specifically supporting the nowadays common hypothesis that lysins from G− usually contain AMP-like regions.

**IMPORTANCE:** Phage-encoded lytic enzymes, also called lysins, are one of the most promising alternatives to common antibiotics. The lysins potential as novel antimicrobials to tackle antibiotic-resistant bacteria not only arises from features such as a lower chance to provoke resistance, but also from their versatility as synthetic biology parts. Functional modules derived from lysins are currently being used for the design of novel antimicrobials with desired properties. This study provides a view of the lysins diversity landscape by examining a set of phage lysin genes. This way, we have uncovered the fundamental differences between the lysins from phages that infect bacteria with different superficial architectures, and, thus, also the reach of their specialization regarding cell wall structures. These results provide clarity and evidences to sustain some of the common hypothesis in current literature, as well as make available an updated and characterized database of lysins sequences for further developments.

## INTRODUCTION

Since the antibiotic pipeline started drying out, a worrying increase in the antibiotic resistant fraction of bacterial populations has been reported (1, 2), and highly antibiotic-resistant population percentages have maintained (3, 4). Thus, if the situation is set to continue, the cost, both economic and in human lives, will be enormous due to the lack of effective treatments (5, 6). This has prompted the interest in novel antimicrobials development by many public health actors, such as international overseeing organizations (7), public health and disease control agencies (3, 4), governments (8), researchers, and several companies (9). Some of the current efforts to gather a new antimicrobial armamentarium have led science towards bacteriophages (phages) (10, 11).

To allow the dissemination of the progeny, double-stranded DNA phages provoke the bacterial host lysis, which is fundamentally accomplished by degradation of the peptidoglycan. This polymer is an essential constituent of the bacterial cell wall, and the breakage of specific bonds within its three-dimensional mesh leads to bacterial death, largely by osmotic shock. The main phage molecule responsible for peptidoglycan degradation is the lysin (also referred to as endolysin). Lysins are released towards their polymeric target, usually with the assistance of another kind of proteins, the holins, which create pores in the plasma membrane and thus allow lysins leakage to the periplasm (12). There are also other phage products that collaborate in hampering the cell wall in some of its particular settings; for example, lysins B detach the arabino-mycolyl outer layer of mycobacteria and their relatives (*e.g., Rhodococcus, Corynebacterium*) (13). Besides, in some Gram-negative bacteria (G−), effective lysis also needs the concurrence of additional phage products named spanins (14). This reveals the important amount of genetic resources put up by phages to overcome the barriers that the bacterial cell walls represent.

In addition to using whole phage particles as therapeutic agents against bacterial infections (the so-called ‘phage therapy’), current efforts also point out to artificially repurposing certain phage products, such as lysins, as antimicrobials (‘enzybiotics’) themselves (11, 15, 16). The concept is rather simple: the external addition of purified lysins to a susceptible bacterium would cause bacterial lysis whenever the lysin degrades the peptidoglycan. This process has been shown to be straightforward in the case of Gram-positive bacteria (G+) and the therapeutic effect of enzybiotics on G+ has been fully confirmed experimentally (15). The most important characteristics that make enzybiotics amenable to be postulated as therapeutics are: a) a certain specificity towards the original bacterial host and some closely related bacteria, that would prevent normal microbiota to be harmed (16, 17), or, conversely, the possibility to have broad-range lysins, if needed (18); b) a lower chance to provoke the appearance of resistant bacteria, which is speculated to be because of the essential nature of the highly conserved peptidoglycan (this is, changes in its structure lead to a decreased fitness and/or virulence) (19); c) neither adverse immune responses nor production of neutralizing antibodies are expected, possibly due to the usual presence of phages —and their products— among the normal cohabitating microbial populations in humans (20). Moreover, lysins are amenable for protein engineering strategies (18, 21–24). Typically, the architectural organization of lysins comprises one (or more) enzymatically active domains (EAD) together with a cell wall-binding one (CWBD). Therefore, synthetic biology strategies, such as construction of completely new lysins made up of different modules as “building blocks”, have been shown to be achievable. Such strategies enable the design and production of tailor-made antimicrobials, based on the conjunction of diverse functions of interest into a single protein. Functions of interest may include, besides a catalytic activity against the peptidoglycan network (*i.e.,* an antimicrobial activity), an increased stability in complex media (25) or, more typically, a certain tropism towards a specific element on the bacterial surface (26) or some other macromolecules like cellulose (27). The engineering approaches mentioned above have circumvented the alleged inability of lysins to cross the outer membrane (OM) of G− (28, 29). Different kinds of synthetic lysins have been devised to that end. Among them we can mention the so-called ‘artilysins’, which are lysins fused to different kinds of membrane permeabilizing peptides (30), the ‘lysocins’, which are lysins fused to elements from bacteriocins that enable bacterial surface recognition and import into the periplasm (22) and the ‘innolysins’, lysins fused to phage receptor-binding proteins (31).

However, a number of lysins also encompass intrinsic bactericidal activity on G− (32–34). This activity was first noticed for the T4 phage lysozyme (35) and several *Pseudomonas aeruginosa* phage lysins (36). Such unexpected property was attributed to non-enzymatic mechanisms, previously described in partially denatured hen egg-white lysozyme (37), and relies on the presence of antimicrobial peptide (AMP)-like subdomains within such lysins, usually at a C-terminal position (32, 38). Recently, it has been suggested that such AMP-like elements are widespread among lysins from phages infecting G−, and that they might cooperate to host lysis by providing an additional affinity towards the cell wall, because of their high net charge (28, 33, 39–41). Since most lysins from G− are assumed to be monomodular, such AMP-like elements are thought to be an alternative to the CWBDs found in multimodular G+ phage lysins for substrate binding. However, it has not been yet properly examined how widespread this trait would actually be, and, therefore, its true functional and evolutionary implications are largely unknown. Of note, such AMP-like elements have been successfully used to design AMPs active on their own (36, 39, 42).

To uncover the actual evolutionary relevance of these AMP-like elements, as well as other lysin features, such as their domain architecture, in this work, a bioinformatic approach examining a wide collection of lysins has been proposed. There are several precedents on the application of homology-based analysis of putative lysin sequences that have paved the way to the systemic comprehension of the co-evolution of phage lysins and their hosts (13, 43). The present study aims to update the picture with the latest available information, as well as to provide answers to the recent questions brought forward by the lysin engineering literature. Therefore, based on current knowledge on the matter and available genomic data, we have constructed and curated a comprehensive database of phage lysin sequences. Subsequent analyses on the data included: a) an initial exploration of the database composition; b) a cross reference of information added to the database to check for differential distribution of distinct domain families and their architectural combinations along different bacterial groups; and c) an overview of easily computable physicochemical properties (net charge, hydrophobicity, etc.) along amino acid (aa) sequences to explore widespread, relevant differences between groups. The hereby conclusions shall, then, strengthen our understanding of lysins specificity and variability, and help in future drug design efforts based on phage products.

## RESULTS AND DISCUSSION

### Outline

A total of 9,539 genomes were prospectively obtained from the National Center for Biotechnology Information (NCBI) database (retrieved on April, 2020). After a careful curation process (for details, see Methods), the final database contained 2,182 proteins and a total of 3,303 Pfam (PF) hits (Table S1 in the supplemental material). Each of these sequences was associated with a bacterial genus corresponding to its described host, for which data on its Gram group and peptidoglycan chemotype was added (Table S2 in the supplemental material). In total, our database comprised phage lysins from 47 bacterial genera, accounting for up to a total of 2,179 sequences, plus three lysin sequences from PRD1-like phages that infect several enterobacteria. Taking into account all of the identical sequences, the 2,182 different sequences of our data set correspond, in fact, to 36,365 entries in the NCBI Reference Sequence database (RefSeq; release 202) (44).

### General differences among lysins

For 1,512 out of 2,182 sequences (69.3%), only one significant PF hit could be predicted (Fig. 1A). This was especially relevant for lysins from phages infecting G−, given that 90.6% of these proteins were predicted to contain a single functional domain. Near 60% of the lysins from phages infecting G+ (for the sake of this work, mycobacteria and their relatives like *Rhodococcus* or *Corynebacterium* were included among G+), harboured only one functional domain. Few lysins appear to contain ≥ 4 PF hits (Fig. 1A). However, these figures should be considered with caution since they do not correspond to the number of real functional modules within the protein, but to a relatively high number (up to 5) of individual repeats that, together, make up a single functional module. For example, the 37 sequences with 6 PF hits correspond to streptococcal phage lysins having the typical structure *[EAD]5×[CW_binding_1]*, being EAD either *Amidase_2* (31 hits), *Glyco_hydro_25* (3 hits) or *CHAP* (3 hits) domains. Likewise, not all sequences with a single PF hit should be assumed to contain only a single domain since many of them might contain other, still undefined domains. Also, some repeats (or even full domains) might not be appropriately predicted if there is enough evolutionary sequence divergence. As an example, the domain structure based upon the three-dimensional folding of pneumococcal major autolysin LytA (45) does not concur with the domains predicted by an homology search since such method is unable to uncover the latest CWBD repeat (Fig. S1 in the supplemental material).

**FIG 1.**
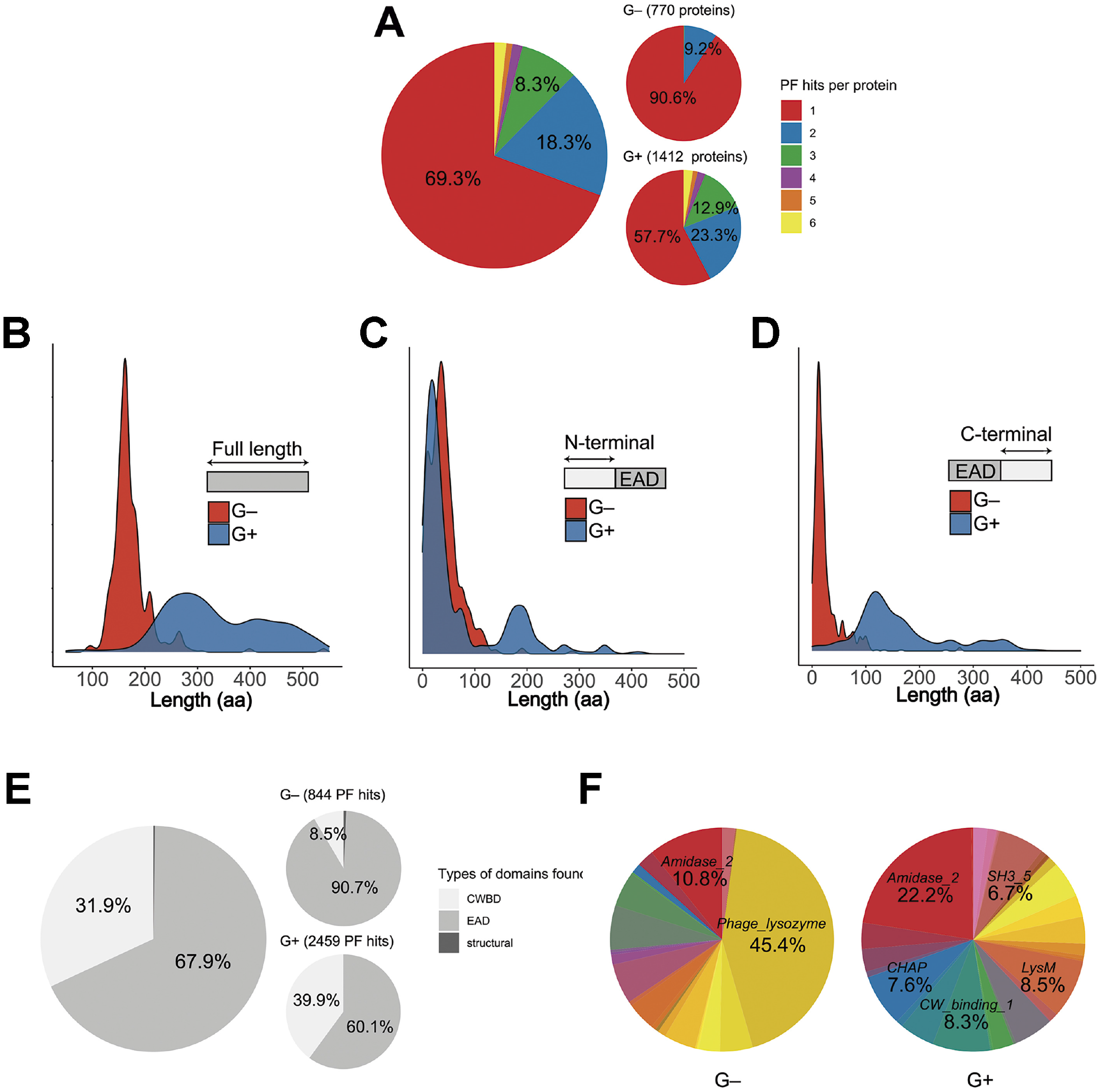
General properties of lysins from phages that infect G+ or G− bacteria. (A) Distribution of the number of PF hits predicted per protein. (B) Distribution of protein lengths. (C and D) Distributions of the number of aa before (C) or after (D) predicted EADs. (E) Distribution of domain types. (F) PF domains variability (different colours stand for different PF domain families, corresponding to those shown in Table 1). In distribution charts (B, C, D) Y-axis shows an estimation of the distribution density.

As a whole, however, the differential relative amount of single and multiple PF hits sequences between G− and G+ phage lysins (Fig. 1A and E) can be taken into account, in accordance with the usual proposal that G− lysins are typically monomodular, while G+ ones are multimodular (46). This is further supported by the evident difference in protein length distributions (Fig. 1B), where G+ phage lysins tend to be larger (median = 317 aa residues) than G− ones (median = 164 aa residues); and also by the differential distribution of sequence lengths before and after the predicted EADs (Fig. 1C and 1D). Fig. 1C shows that EADs from G− phage lysins start, approximately, at the same point than G+ ones, this is, near to the N-terminal end of the protein, except that the EADs starting point distribution is slightly shifted towards the C-terminal part of the enzyme in lysins from G−, probably due to the presence, in some cases, of CWBDs at the N-terminus (28). Of note, G+ EADs starting point distribution shows a secondary local maximum at around coordinate 200. This is consistent with the presence of EADs at a medial location within the protein, something that has already been observed in many G+ phage lysins (13, 47). According to Fig. 1D, most G+ EAD hits have much more “space” at the C-terminal part than G− ones (respective medians of C-terminal length after EAD hit distributions for G− and G+ are 16 and 136 aa residues). The additional length at the C-terminal part of G+ phage lysins must be occupied by non-catalytic domains (*i.e.,* CWBDs) and, taken together, all this evidence would support the common postulate that most detected G− lysins are monomodular.

Finally, Fig. 1F illustrates that, in contrast with the case of G− lysins, G+ lysins present a high diversity of different types of domains. There is a remarkable predominance of the EADs belonging to the *Phage_lysozyme* family of proteins in G− lysins (45.4% of total hits), whereas *Amidase_2*, the most frequent EAD among G+ phage lysins, accounted only for 22.2% of G+ PF hits.

### Differential distribution of domain families among different bacterial host groups

A distribution analysis of each PF family amongst bacterial hosts was performed (Table 1). From the total 3,303 PF hits analysed, 2,460 corresponded to phages infecting G+ bacteria. 2,243 (1,477 G+; 766 G−), 1,054 (982 G+; 72 G−), and 6 (G−) corresponded to EADs, CWBDs, and structural domains, respectively (the sources for domains classification as EAD, CWBD or structural, can be consulted at Table S3 in the supplemental material). When the differential Gram group classification of each PF hit was analyzed, it was found that EADs like *Amidase_5, Glyco_hydro_25, Peptidase_C39_2*, and *Transglycosylase* were exclusive of G+, whereas *Glyco_hydro_108* or *Muramidase* were characteristic of phages infecting G−. Other EADs like *Amidase_2, Amidase_3, CHAP, Glucosaminidase*, *Peptidase_M15_4*, and *Peptidase_M23* were common in G+, whereas *Glyco_hydro_19, Hydrolase_2, Phage_lysozyme* dominated amongst G−. Besides, *CW_7, CW_binding_1, LGFP, SH3_5*, or *ZoocinA_TRD* constituted the CWBDs of G+, and, although *LysM* and *PG_binding_1* were most frequently found in G+ lysins, also appeared sometimes among G− (Table 1 and Fig. 2). *PG_binding_3* was the only CWBD exclusive of G− lysins. Interestingly, all of the 40 *PG_binding_3* occurrences were accompanied by *Glyco_hydro_108* at the N-terminal moiety, yielding an architecture *([Glyco_hydro_108][PG_binding_3])* that was widespread among γ-proteobacteria.

**Table 1.** Distribution of PF hits of phage lysins from Gram-positive and Gram-negative bacteria^*a*^

| Domain | Domain type | G+ (%) | G− (%) | Total | Encoded by phages of: |
| --- | --- | --- | --- | --- | --- |
| *3D* | EAD | 7 |  | 7 |  |
| *Amidase_2* | EAD | 547 (85.7) | 91 (14.3) | 638 | Widely distributed in G+ |
| *Amidase_3* | EAD | 93 (81.6) | 21 (1.4) | 114 | *Bacillus*, *Streptococcus*,<br>*Clostridium*, *Staphylococcus* |
| *Amidase_5* | EAD | 81 (100) |  | 81 | *Streptococcus*, *Lactococcus* |
| *CHAP* | EAD | 186 (93.9) | 12 (6.1) | 198 | *Streptococcus*,<br>*Staphylococcus*,<br>*Streptomyces*, *Arthrobacter* |
| *Cutinase* | EAD | 24 |  | 24 |  |
| *FSH1* | EAD | 3 |  | 3 |  |
| *Glucosaminidase* | EAD | 73 (97.3) | 2 (2.7) | 75 | *Streptococcus* |
| *Glyco_hydro_19* | EAD | 9 (14.3) | 54 (85.7) | 63 | *Acinetobacter* and other<br>genera |
| *Glyco_hydro_25* | EAD | 142 (100) |  | 142 | *Lactobacillus*, *Bacillus*,<br>*Streptococcus* |
| *Glyco_hydro_108* | EAD |  | 43 (100) | 43 | Widely distributed in G− |
| *GPW_gp25* | EAD |  | 12 | 12 |  |
| *Hydrolase_2* | EAD | 3 (5.8) | 49 (94.2) | 52 | *Escherichia*, *Pseudomonas*,<br>*Vibrio* |
| *Muramidase* | EAD |  | 35 (100) | 35 | *Pseudomonas*, *Burkholderia* |
| *NLPC_P60* | EAD | 7 | 6 | 13 |  |
| *PE-PPE* | EAD | 13 |  | 13 |  |
| *Peptidase_C39_2* | EAD | 42 (100) |  | 42 | *Mycobacterium* |
| *Peptidase_C93* | EAD |  | 4 | 4 |  |
| *Peptidase_M15_3* | EAD |  | 10 | 10 |  |
| *Peptidase_M15_4* | EAD | 96 (71.1) | 39 (28.9) | 135 | *Mycobacterium, Bacillus* |
| *Peptidase_M23* | EAD | 77 (97.5) | 2 (2.5) | 79 | *Arthrobacter, Mycobacterium, Rhodococcus* |
| *Pesticin* | EAD |  | 2 | 2 |  |
| *Phage_lysozyme* | EAD | 32 (8.0) | 366 (92.0) | 398 | Widely distributed in G– |
| *Phage_lysozyme2* | EAD | 1 |  | 1 |  |
| *Prok-JAB* | EAD |  | 1 | 1 |  |
| *Prophage_tail* | EAD | 3 |  | 3 |  |
| *SLT* | EAD | 6 | 16 | 22 |  |
| *Transglycosylase* | EAD | 32 (100) |  | 32 | *Mycobacterium* |
| *Amidase02_C* | CWBD | 20 |  | 20 |  |
| *Big_2* | CWBD | 1 |  | 1 |  |
| *CW_7* | CWBD | 125 (100) |  | 125 | *Streptococcus, Arthrobacter, Streptomyces* |
| *CW_binding_1* | CWBD (repeat) | 205 (100) |  | 205 | *Streptococcus* |
| *CW_binding_2* | CWBD (repeat) | 3 |  | 3 |  |
| *DUF3597* | CWBD | 5 |  | 5 |  |
| *LGFP* | CWBD (repeat) | 31 (100) |  | 31 | *Rhodococcus* |
| *LysM* | CWBD | 210 (97.7) | 5 (2.3) | 215 | Widely distributed in G+ |
| *PG_binding_1* | CWBD | 130 (83.3) | 26 (16.7) | 156 | *Mycobacterium, Bacillus, Streptomyces* |
| *PG_binding_3* | CWBD |  | 40 (100) | 40 | Widely distributed in G– |
| *PSA_CBD* | CWBD | 13 |  | 13 |  |
| *SH3_3* | CWBD | 20 | 1 | 21 |  |
| *SH3_5* | CWBD | 164 (100) |  | 164 | *Streptococcus, Staphylococcus, Lactobacillus, Bacillus* |
| *SPOR* | CWBD | 5 |  | 5 |  |
| *ZoocinA_TRD* | CWBD | 50 (100) |  | 50 | *Streptococcus, Enterococcus* |
| *Gp5_C* | Structural |  | 3 | 3 |  |
| *Gp5_OB* | Structural |  | 3 | 3 |  |

<sup>a</sup> Percentages and further remarks are only shown for domains represented by, at least, 30 hits

**FIG 2.**
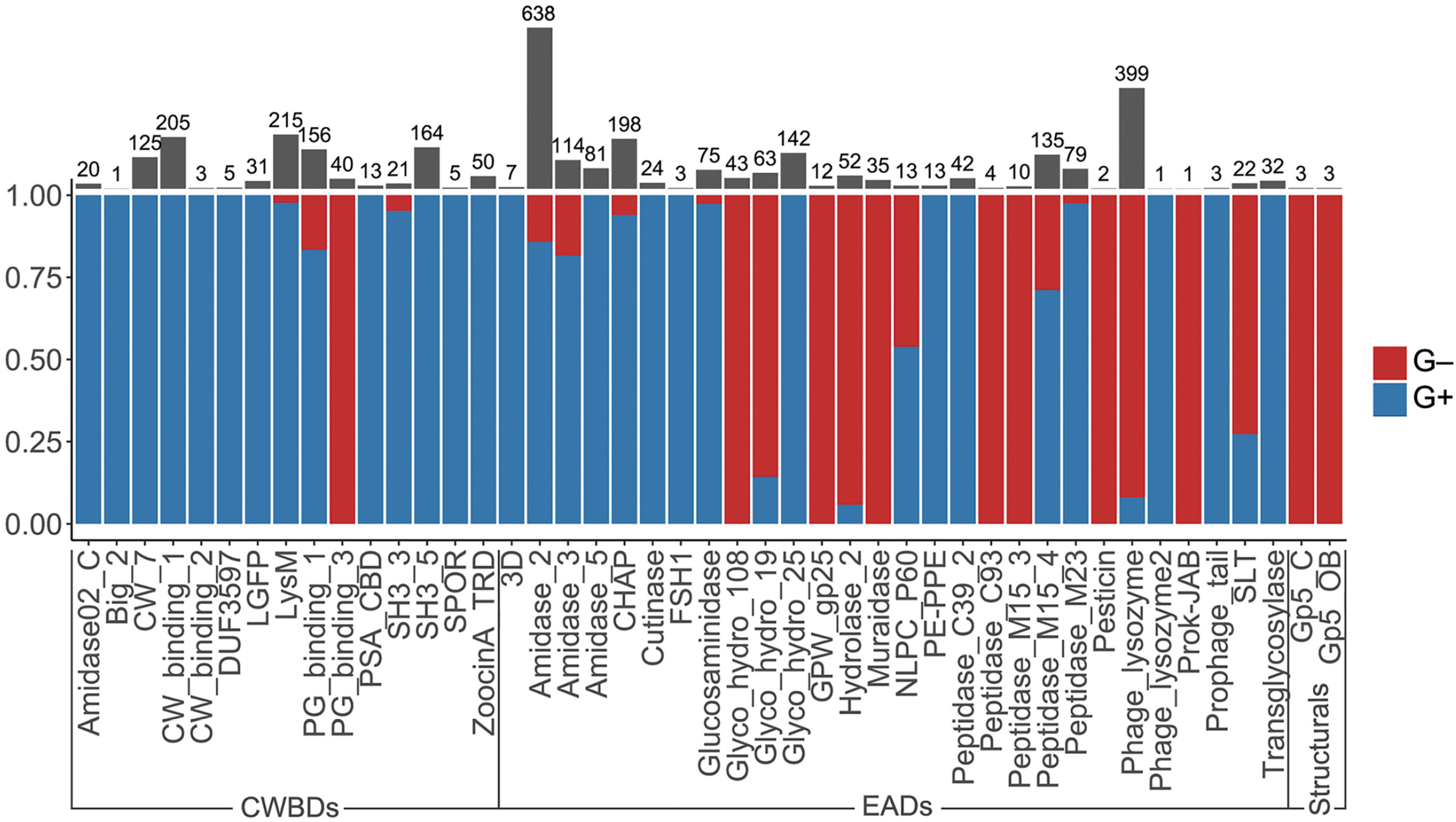
Differential distribution of PF hits among G− and G+ bacterial hosts. Y-axis shows the proportion of PF hits found in G+ within a given domain family. Grey bars and numbers above represent the total number of hits of each PF domain.

Trends in PF domains distribution among genera, rather than Gram group, were a bit more complex (Figs. 3 and 4), although some conclusions could be reached. To begin with G+ CWBDs, the *CW_binding_1* repeats were only encoded by phages infecting streptococci, whereas *CW_7* constitute the CWBD of many phage lysins of *Streptococcus, Arthrobacter*, and *Streptomyces*. *CW_binding_1* repeats are known to bind choline residues present in the teichoic acids of *Streptococcus pneumoniae* and its relatives (*i.e.,* streptococci of the Mitis group) (48, 49), and therefore only appeared within our dataset among such group of bacterial hosts (Fig. S2 in the supplemental material). CW_7 repeats are known to bind a conserved peptidoglycan motif, and are thus less restricted in the variety of bacteria they may recognize (50). *LysM* domains were also widely distributed in G+, *ZoocinA_TRD* was very common among *Streptococcus thermophilus* and *PSA_CBD* was exclusive for *Listeria* phage lysins. As for EADs, *Amidase_5* was very frequently found among streptococci and *Amidase_2* generally abundant among all G+.

**FIG 3.**
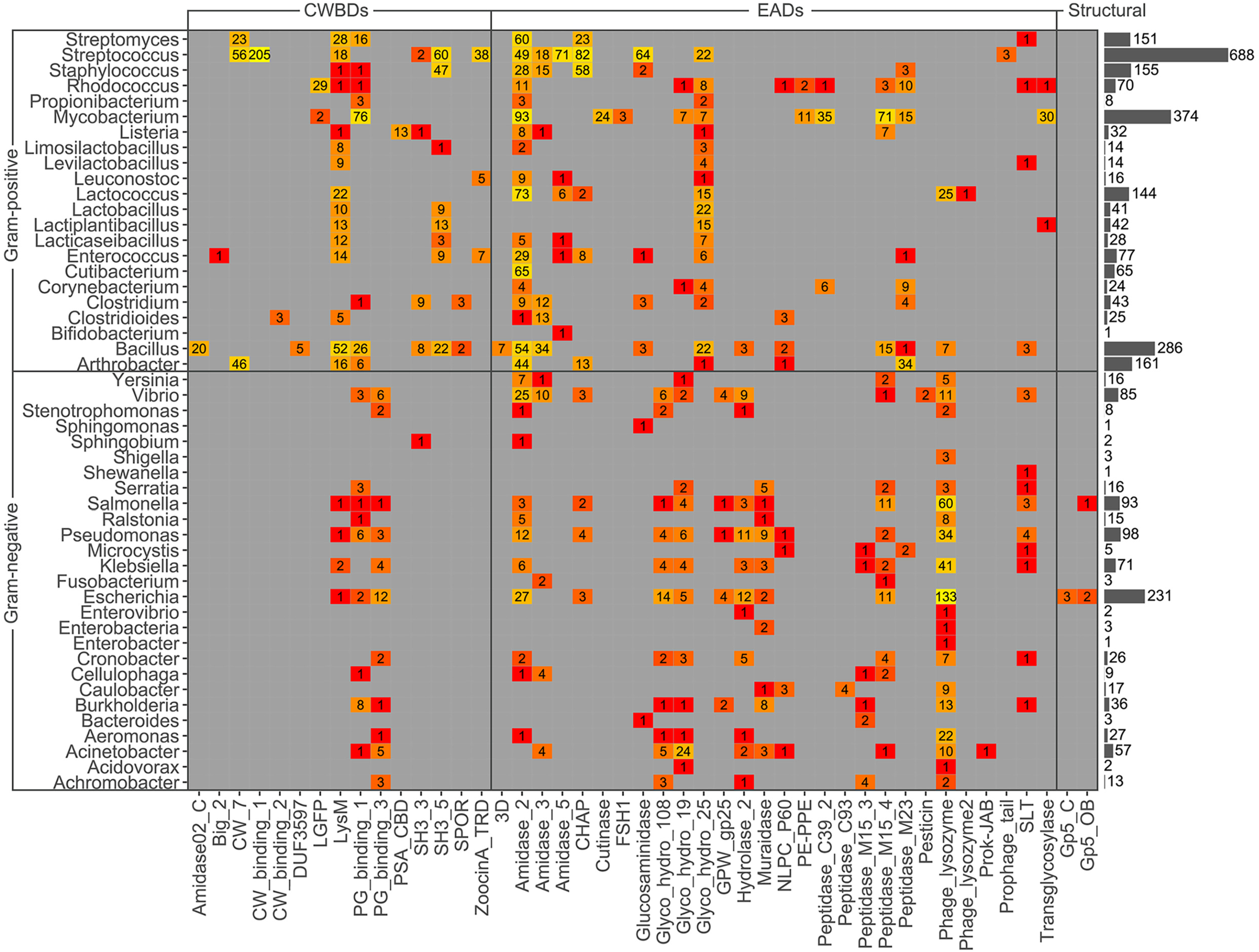
Heatmap of PF hits distribution across host bacterium genera. Numbers within each tile indicate the number of hits predicted for the corresponding taxon and PF family. The colour scale represents the number of hits from low (red) to high number (yellow). Grey bars at the right represent the total number of PF hits predicted within each genus.

**FIG 4.**
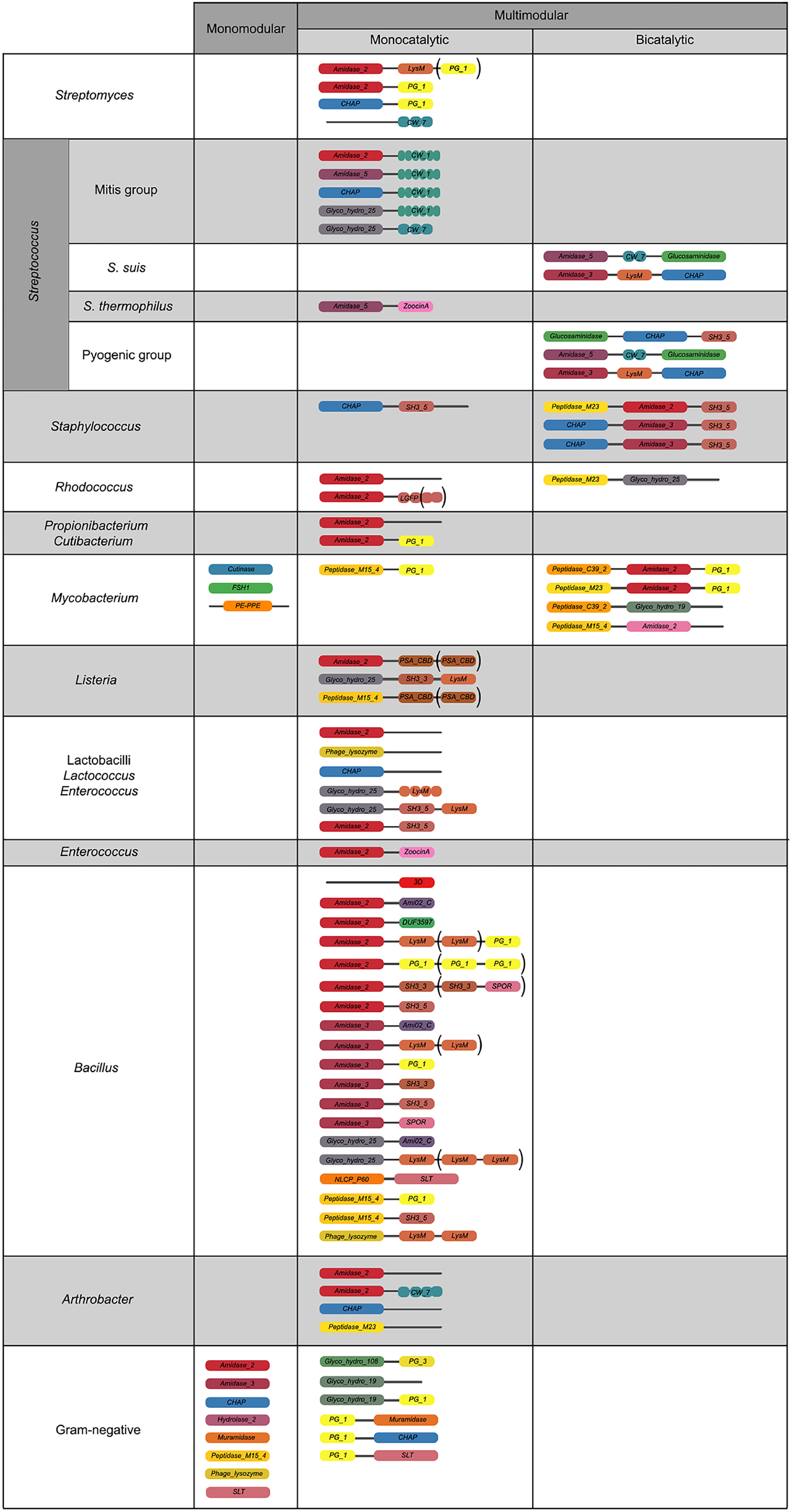
Relevant architectures observed in lysins from phages infecting different taxonomic groups of bacteria. Different colours mean different domains; brackets denote domains that appear only in some representatives of the depicted architecture.

Another exclusive trait of some G+ lysins was the concurrence of two distinct EADs. This was observed for phage lysins from *Streptococcus suis*, Pyogenic group streptococci, staphylococci or mycobacteria. A possible explanation for multicatalytic lysins is an increased lytic efficiency over monocatalytic ones, since activities attacking different sites of the peptidoglycan are known to act synergistically in peptidoglycan degradation (51). Such synergy could also imply a decreased chance for the appearance of resistant peptidoglycan mutants (52). It has also been shown that the synergistic concurrence of both activities is sometimes needed for full activity. Thus, it has been suggested that some phages may have evolved a regulatory mechanism to avoid lysis of other potential host cells relying on the proteolysis of bicatalytic lysins by host-cell proteases. Then, both EADs would be disjointed by proteolysis upon host cell lysis and the degraded lysins would no longer be active against the nearby bacterial population (53). This should be especially relevant for phages infecting G+ bacteria, which lack a protective OM hindering the lysis of other bacterial cells from without, and hence the exclusiveness of the bicatalytic architecture among phages infecting G+. In some other cases, however, it is the high affinity of the CWBD that has been proposed as the mechanism that maintains lysins tightly bound to cell debris preventing widespread lysis of the bacterial community (54), which is also an argument for the widespread presence of CWBDs among G+ and not among G−.

Staphylococcal phage lysins presented reduced EAD variability, normally using *Amidase_2, Amidase_3* and/or *CHAP* domains, with *SH3B_5* being the preferred CWBD, in agreement with previous results (55). In some cases, the staphylococcal *SH3B_5* has been shown to bind the peptidoglycan with the characteristic pentaglycine interpeptidic bridge of *Staphylococcus* (56). Domains putatively assigned an esterase activity *(Cutinase, FSH1, PE-PPE)* were only present in phages from *Mycobacterium* and its relatives, presumably as type B lysins. The *LGFP* repeats, quite common among *Rhodococcus* phages, might be a specific CWBD among such Corynebacteriales. Peptidase EADs were common and diverse among mycobacteriophages, in contrast with other G+ phages, which do not typically contain peptidase EADs other than *CHAP*. Of note, *CHAP* domains have been sometimes described as peptidases but, in other occasions, as *N*-acetylmuramoyl-L-alanine amidases (NAM-amidases) (57, 58).

Regarding G−, the most widely spread architecture of G− phage lysins was monomodular, harbouring a single *Phage_lysozyme* domain, which accounted for half (50.8%) of the identified G− lysins in our database. Another architecture that was only found in G− lysins is the localization of a CWBD at N-terminal end (for example, as *[PG binding l] [Muramidase])*, although they were not at such position in every case *(e.g.*, architecture *[Glyco_hydro_108] [PG_binding_3]* was also present).

The correlation between domain distribution and peptidoglycan composition might also shed some light on the relationships of different domain families with different taxa. To that end, the chemotypes classification of peptidoglycan proposed by Schleifer and Kandler (59) was used (Table S2 in the supplemental material). Briefly, such classification hierarchically relies on (i) the site of cross-linkage of the peptide subunit of the peptidoglycan, (ii) the nature of the cross-link and (iii) the specific residue at position 3 within such peptide subunit (Fig. 5A). Starting by CWBDs (Fig. 5B), classification by chemotypes did not provide a better explanation for specificity than other genera-specific traits, as discussed above. Some specificities could be found though (*e.g*., *Amidase02_C* appears only in phages that infect A1α bacteria or *PG_binding_3* only in A1γ), and some CWBDs that are widespread among different chemotypes could also be observed *(PG_binding_1, LysM, SH3_5)*. In general, however, it cannot be stated that peptidoglycan composition is a major determinant for CWBD specificity, except for some cases such as, for example, *ZoocinA_TRD* domains, which has been proposed to bind A3α with two Ala residues at the cross-link (60). The poor performance of chemotype as an *a priori* predictor of the CWBD PF family ligand is more clearly evident if we consider the CWBD types which appeared widespread among many different chemotypes, such as *LysM* and *SH3_5*. To check whether this apparent ‘promiscuity’ may be linked to the presence of subfamilies with potentially different ligands or if it could rather be a true promiscuous binding, SSNs were constructed with the PF hits of *LysM* and *SH3_5* (Fig. S3 in the supplemental material). The *LysM* SSNs did not show prominent similarity clusters either classified by taxon or by chemotype of the bacterial host. This suggests that *LysM* could be a truly ‘universal’ CWBD that would bind to a conserved cell wall ligand. The rather generic description of *LysM* ligands in the literature (as ‘N-acetylglucosamine-containing polysaccharides’) is in agreement with this observation. *SH3_5*, however, displayed at least two differentiated sequence similarity groups that correlated rather well with different taxonomic groups (namely, staphylococci versus streptococci and lactobacilli). In fact, literature reflects that, while lytic enzymes with predicted *SH3_5* domains typically recognize polysaccharides (and peptidoglycan in particular), there seem to be different specializations. For example, the CWBD of the *Lactiplantibacillus plantarum* major autolysin binds many different peptidoglycans with low affinity, being glucosamine the minimal binding motif (61), while *SH3_5* domains from staphylolytic enzymes have been shown to be rather specific to crosslinked peptidoglycans (like the A3α peptidoglycan of *Staphylococcus* and *Streptococcus*) and that the nature of the crosslink itself determines the affinity of such CWBDs for the peptidoglycan (62, 63).

**FIG 5.**
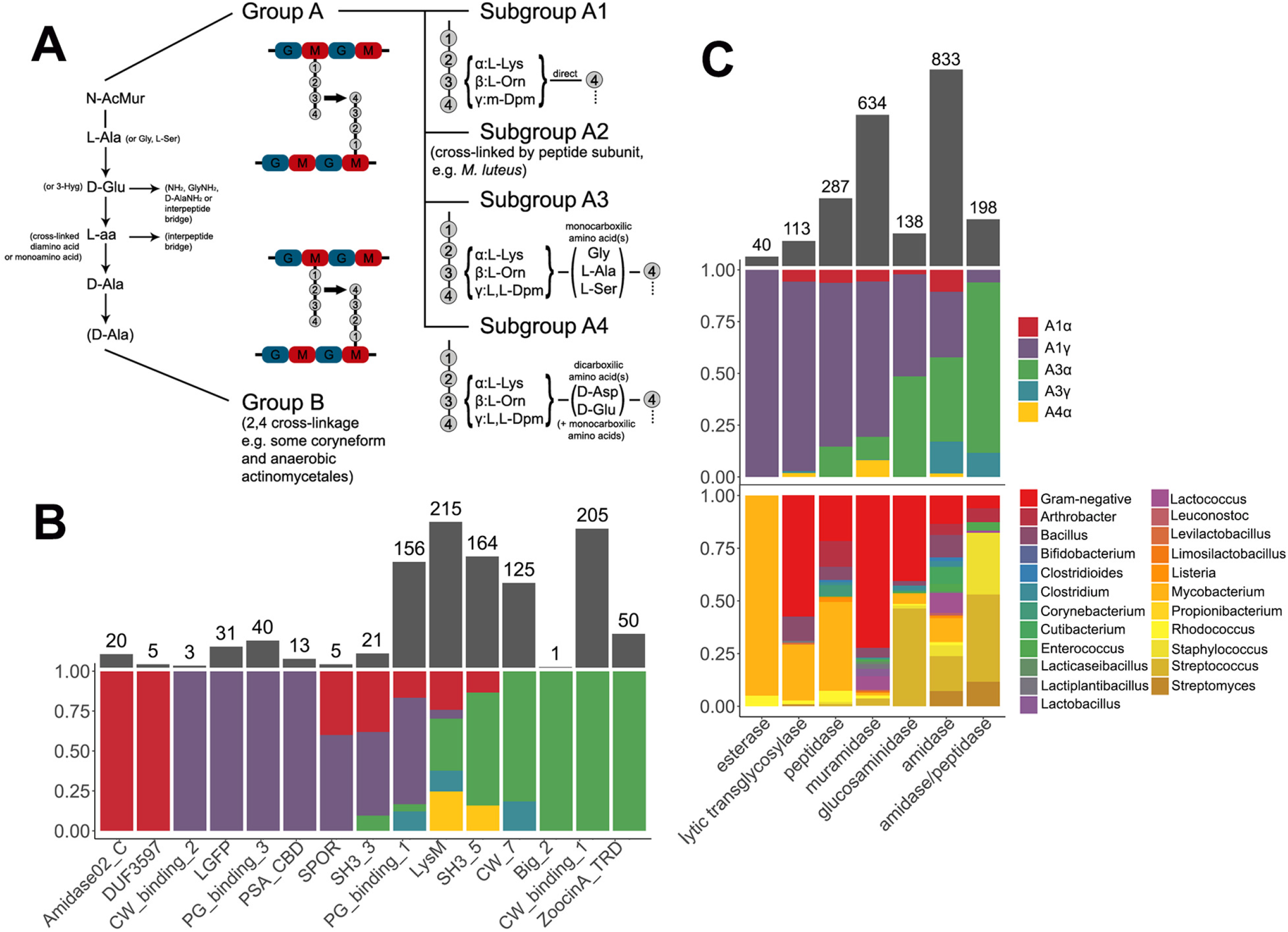
Differential distribution of CWBDs and catalytic activities across peptidoglycan chemotypes and taxonomic groups of bacterial hosts. (A) Schematic representation of the relevant peptidoglycan chemotypes present for the bacterial hosts in our dataset. (B) Distribution of CWBD PF hits among chemotypes. (C) Distribution of catalytic activities of EAD PF hits among chemotypes and taxonomic groups.

Additional information could be drawn from this analysis when applied to the different catalytic activities detected (Fig. 5C). First of all, NAM-amidases were the most represented type of domains and also those that appeared among more different taxonomic groups and chemotypes, even more so than lysozymes. Indeed, *Amidase_2*, the most abundant PF domain in our dataset (638 hits), appeared both in lysins from G+ and G− phages. The SSN in Fig. S3 in the supplemental material shows, however, that although *Amidase_2* seems a rather diverse group, with various observable similarity clusters, none of such clusters correlate with any of the classifiers of the bacterial hosts tested.

Muramidases were quite overrepresented among G− bacteria (chemotype A1γ) because of the widespread presence of *Phage_lysozyme* domains. Glucosaminidases appeared evenly both against A1 and A3 peptidoglycans, but whereas in G+ bacteria (which comprise all A3s and a few A1s) glucosaminidase activity was represented by *Glucosaminidase* PF domain, the only domain putatively assigned with a glucosaminidase activity among G− was *Glyco_hydro_19* (Figs. 3 and 5).

Another interesting remark is that peptidase activities were more common amongst lysins from phages infecting bacteria with subgroup A1 peptidoglycans which, in turn, display the simplest cross-linkage of all types, lacking an interpeptide bridge. Thus, peptidases were not uncommon among G−, and were also present in A1 phages from G+ (especially mycobacteriophages, but also listeriophages and phages from *Clostridium, Bacillus* or *Corynebacterium*). On the other hand, amidase/peptidases, which is the label given to *CHAP* domains (Table S3 in the supplemental material), were much more prevalent among A3 G+, and only seldom present in lysins from phages infecting A1 bacteria (namely some G−). This suggests that if there was to be an A3-specific peptidase activity would be that located in *CHAP* domains. It makes sense that different peptidase structures have evolved towards A1 and A3 peptidoglycans, since the complexity of their peptidoglycan peptide moieties differs significantly. Adding to this conclusion, the *CHAP* SSN (Fig. S3 in the supplemental material) did show a similarity clustering of the few *CHAP* examples in lysins from A1 phages, besides an apparent differentiation of *Staphylococcus* and *Streptococcus*/*Enterococcus*.

### Physicochemical analysis of phage lysins from Gram-positive *versus* those from Gram-negative bacteria

The results analysed so far support a distinct distribution of domain architectures and families among lysins that infect different kinds of bacteria, and even hint to an association of such differential distribution to some cell wall properties. To check whether such variations can also be correlated with a measurable difference in physicochemical properties, net charge, net charge per residue (NCPR), hydrophobicity, average hydrophobic moment, and aliphatic index were calculated and used to implement a random forest (Fig. 6). This way, the aforementioned physicochemical variables were used as classifiers for the prediction of the host bacterium Gram group of lysins. The resulting algorithm yielded a Receiver Operating Characteristic (ROC) plot with an area under the curve (AUC) of 0.897, which can be interpreted as a good predictive ability (Fig. 6A). Using the probability threshold (0.591) derived from the best point of the ROC curve (which maximizes true positive rate and minimizes false positive rate), G+/G− classification upon the testing subset (Fig. 6B) managed an accuracy of 87.9% with sensitivity and specificity, respectively, of 84.1% and 81.3% (being the classification as G+ the “positive” one). According to the subsequent analysis (Fig. 6C), NCPR was the most relevant variable to distinguish between G+ and G−, followed by average hydrophobic moment and aliphatic index and, finally, hydrophobicity. In general, these results suggest that lysins from phages that infect G+ and G− can in fact be differentiated by their physicochemical properties in a relatively efficient manner. For visualization of the differences between G+ and G−, a multidimensional scaling (MDS) plot based on the proximity matrix from the random forest model was drawn (Fig. 6D). Such plot showed the clustering of G− lysins within the 2-dimensional space based on the physicochemical variables, while G+ ones seemed to be more dispersed. A qualitative interpretation of this result may reflect the aforementioned wide diversity of functional modules and architectures of G+ lysins (Fig. 1D, Fig. 4) in contrast with the relatively low variability of G−. Such low variability of the G− lysins hereby analysed would then be associated with a preference for some physicochemical features. The sense of this preference was subsequently checked.

**FIG 6.**
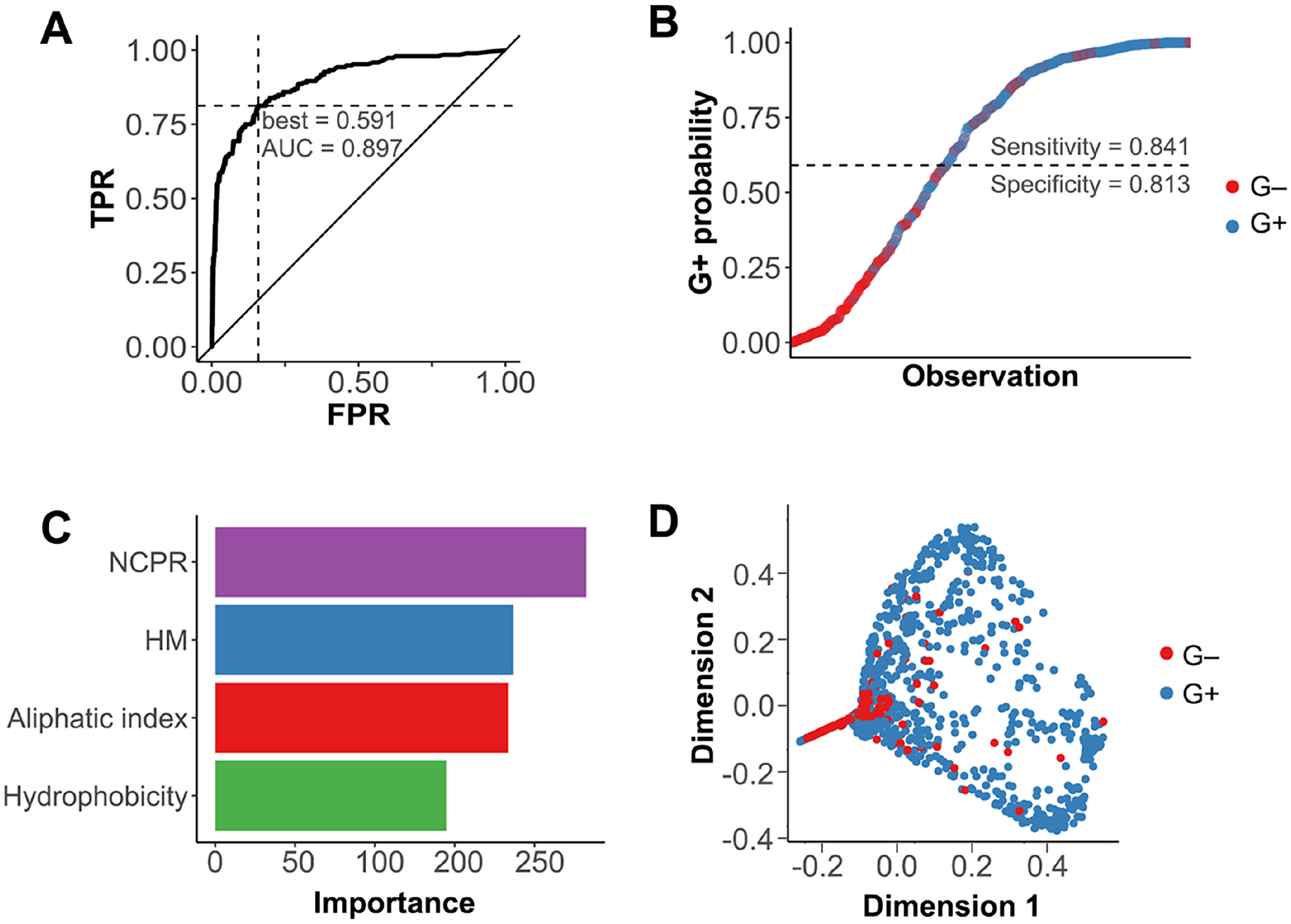
Random forest prediction and classification of Gram group of bacterial host based of lysins physicochemical properties. (A) ROC curve of the random forest predictive model (TRP: true positive rate, FPR: false positive rate). ROC best point of positive group (G+) probability for outcome maximization is presented, as well as the AUC. (B) Random forest casting of bacterial host Gram group on the testing subset of lysin sequences. The dashed line represents the G+ probability threshold for classification based on the ROC best point. (C) Relative importance of each of the four descriptors used for classification within the model. (D) MDS plot of the training subset according to the proximity matrix derived from the random forest.

Indeed, the net charge distribution (normalized by protein length) was significantly higher in G− lysins than in G+ ones (*p* ≤ 0.0001; ES = 0.66) (Fig. 7A, most left panel). Moreover, the average prediction of local net charge suggested that such difference is mainly located at the C-terminal part of G− lysins (Fig. 7B). A more thorough comparison (Fig. 7C) seemed to confirm this. At every sequence quartile of the proteins (*i.e.,* contiguous fragments of sequence with a length equal to 1/4 of the total number of aa residues in the original protein sequence), the net charge distribution of G− lysins had a significantly higher net charge. However, the actual size of this shift was only moderate along the sequences (ES between 0.24 and 0.34) but it was, again, higher at the final quartile (0.52).

**FIG 7.**
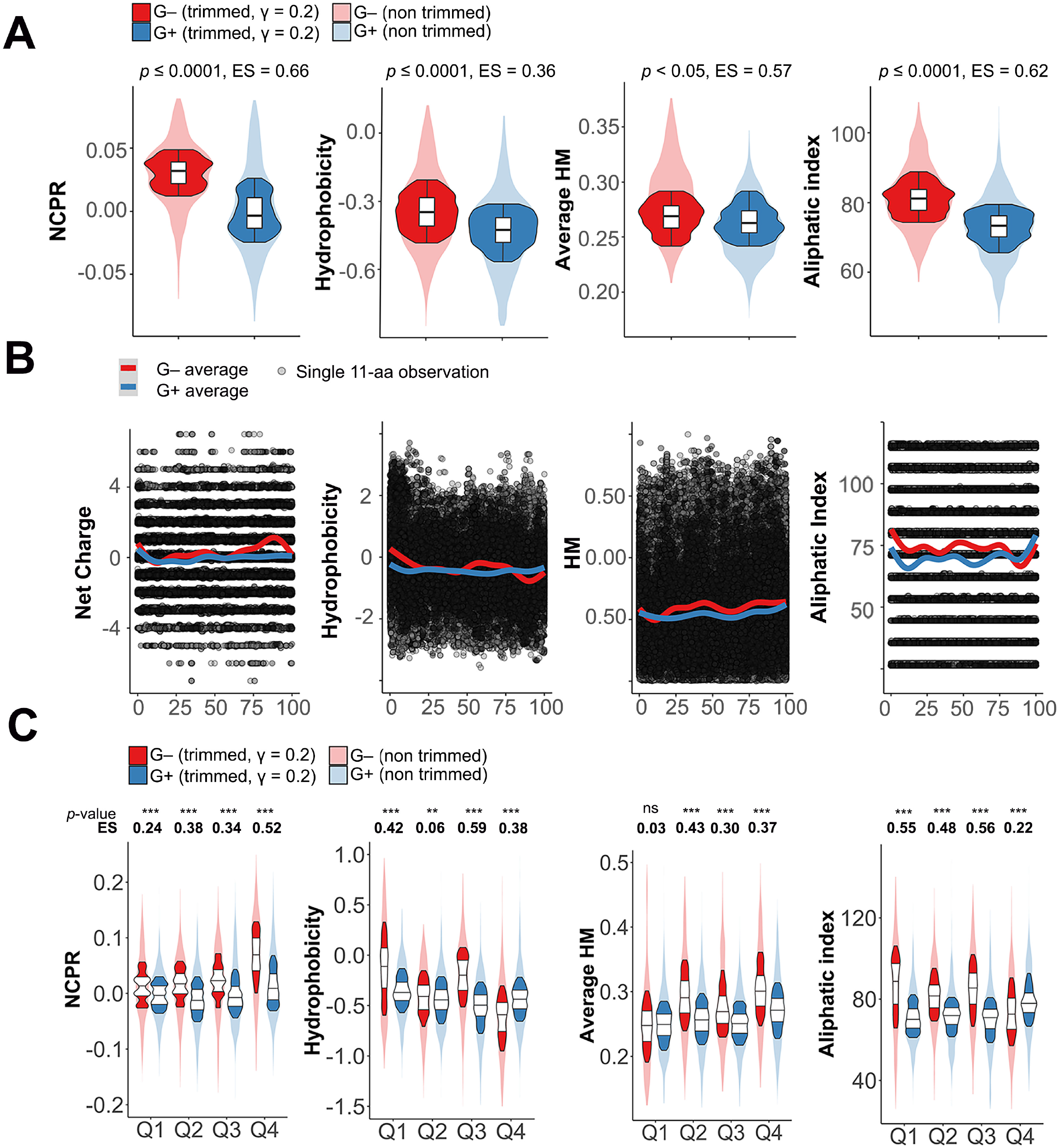
Differential physicochemical properties distribution among G+ and G− phage lysins. (A) Distribution of net properties calculated along the whole protein sequences of lysins from phages infecting G− or G+. (B) Local computation of physicochemical properties. Each dot represents the particular value calculated for an 11-aa window in a given lysin. Continuous lines are average tendencies based on either all G− or all G+ data points. (C) Distribution of different properties at quartiles of lysin sequences. Asterisks indicate p-values (** ≤ 0.01, *** ≤ 0.001) obtained from the Yuen-Welch test for trimmed means with a trimming level of γ = 0.2; ES indicates the Wilcox and Tian’s ζ measurement of effect size.

Hydrophobicity was also higher in G− lysins, but the difference regarding G+ ones is smaller (ζ = 0.36). This might be related to the rather inconsistent pattern shown by average local hydrophobicity and sequence quartiles comparison (Fig. 7BC). G− lysins tended to have a more hydrophobic N-terminal part, whereas at the C-terminal moiety the tendency was reversed, something that can be explained by the relative abundance of positively charged residues shown before for G−. It is at the third quartile (Q3), immediately before the high positive net charge patch described above, where the difference was statistically more relevant (*p* ≤ 0.001; ES = 0.59), with higher values in G−s. There was also a statistically significant difference in the average hydrophobic moment distributions between G+ and G− phage lysins. For the G− group, the local plot (Fig. 7B) showed a higher tendency to present greater hydrophobic moments along the whole protein length but the N-terminal part. Analysis of sequence quartiles confirmed a statistically significant superiority of average hydrophobic moment for G− except at N-terminal. The aliphatic index was also significantly higher in G−, although G+ showed an aliphatic index peak at their C-terminal part that surpassed that of G− (coincidental with G− basic aa peak, which, understandably, would lower both hydrophobicity and aliphatic index at Q4) (Fig. 7C).

Taking all these observations together with the results thrown by the random forest prediction, we can conclude that the physicochemical difference between lysins from phages that infect G+ or G− bacteria is specified as a higher positive net charge of G−, particularly at C-terminal end, combined with a greater propensity in incorporating aliphatic aa and likely resulting in amphiphilic structures.

A closer examination of net charge (and C-terminal net charge) of lysins from G− infecting phages indicated that the high positive patch trait seems specific to some domain families. As a whole, a statistically significant higher NPRC value was found in lysins bearing *Phage_lysozyme, Hydrolase_2* and *Glyco_hydro_19* domain families (Fig. 8). At the C-terminal part, higher NCPR was found in lysins bearing the same domains mentioned above, but also in *SLT* and *Muramidase*. The average local net charge tendency showed for each EAD group (Fig. S4 in the supplemental material) confirmed that a local high positive charge peak appears in the protein part immediately before the C-terminal apex.

**FIG 8.**
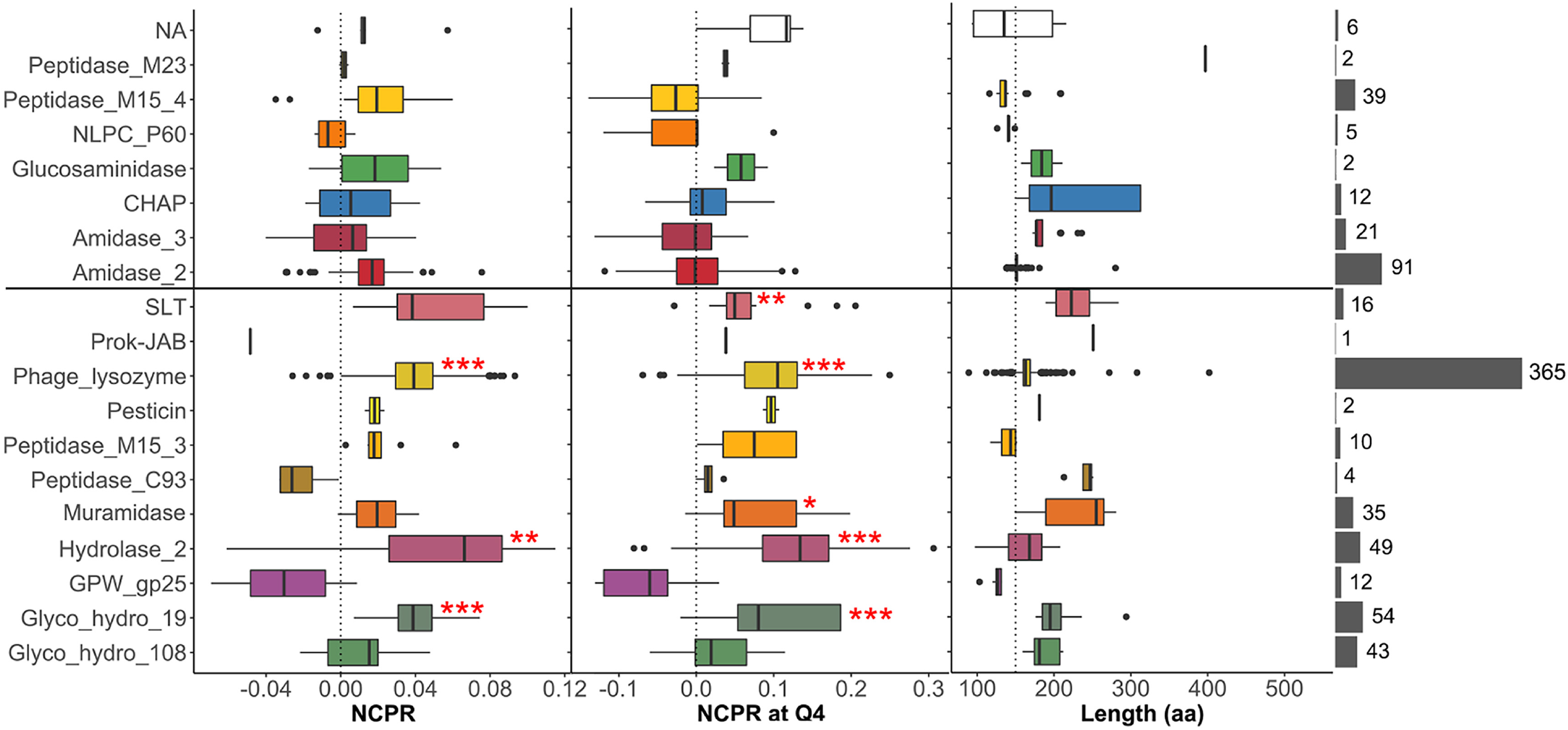
Net charge distribution of lysins from G− infecting phages classified according to the predicted EAD. Rightmost grey bars depict the number of lysins classified into each EAD group (lysins within NA group are those for which an EAD was not assigned). All groups were compared with the distribution of the *Amidase_2* domain, as a highly represented, near-neutral control using Welch’s test on γ = 0.2 trimmed means with *post hoc* Bonferroni correction (*, *p*-value ≤ 0.05; **, *p*-value ≤ 0.01; ***, *p*-value ≤ 0.001).

Interestingly, all of the aforementioned domains that present a higher, positive charge patches at their C-terminal part were preferentially present in lysins from phages that infect G− bacteria (Table 1). This observation provides a basis to argue a generalized evolutionary tendency in G− infecting phages towards developing AMP-like subdomains at the C-terminal moiety of their lysins. Such subdomains contain, indeed, features typical of AMPs (such as the high net charge accompanied by a high local hydrophobic moment, hydrophobic patches, etc.), and may play a role in the interaction between lysins and cell wall in G− bacteria. Electrostatic interactions do play a significant role in phage-bacteria interplay, as suggested for modular lysins from phages that infect G+ bacteria. For example, it has been shown that the negative net charge of many G+ lysins hinders their ability to approach the negatively charged cell wall (18, 64). This renders the affinity-based interaction of the CWBDs with their cell wall ligands essential for lysins activity. This essentiality of CWBDs has been shown for several lysins (20), but generalizations should be made with caution because there are also cases reported of single catalytic domains that lysed G+ cells more efficiently when their CWBD was removed (65). To our knowledge there are only few cases reported in which CWBDs appear to increase the efficiency of cell wall-lysin interaction in G− lysins (21). We have already shown, based on our own data, that it is safe to say that G− lysins are monomodular. Thus, taking this theoretical framework into account, it could be argued that G− lysins should have evolved a distinct strategy to grant cell wall interaction, namely an increased net charge and, perhaps, the presence of hydrophobic patches near such basic residues (*i.e.,* AMP-like regions), rather than containing an additional CWBD, which, incidentally, might be essential for post-lytic regulation in G+, but not in G− (54). The AMP-like subdomains, besides providing better anchorage to bacterial surface structures, might as well act as an additional mechanism towards effective lysis of G− bacteria. There are indeed abundant examples in literature on the ability of G− lysins to interact with the OM and permeabilize it (33, 38, 39, 66), a trait that, it is plausible to say both from our own analysis and the experimental results of many works, would reside in such AMP-like elements. If we assume this, the identification of AMP-like subdomains within lysins could provide also a way of predicting the ability of such lysins to better interact with the OM from without, and thus their antimicrobial potential.

### Concluding remarks

Phages and their bacterial hosts are constantly evolving in a co-dependent manner (67). From the point of view of phage lysins, this means that such molecules have adapted to the particular structures and features of the host cells. This adaptation can be described as the functional adjustment of the protein elements to optimally fulfil their purposes: the efficient and regulated degradation of the peptidoglycan. Therefore, lysin structures and cell wall structures must be closely correlated. A way of testing and understanding such relationship was the hereby presented sequence-based classification of the domains constituent of phage lysins, and the analysis of their distribution among (pseudo)taxonomical and structural classes of bacterial hosts. Our procedure yielded several important associations of lysins and cell wall architectures explainable in a structural-functional way:

a. The different architectures found between lysins from phages that infect G+ or G−. The ones from G− are usually monomodular, whereas lysins from G+ infecting phages are multimodular. Moreover, the bicatalytic type of modular structure only appears among G+. An explanation for this architecture is the requirement for a tighter post-lytic regulation in G+ and/or a more efficient lytic activity relying on a tighter substrate binding or on the synergistic effect of combining different catalytic activities.
b. The association of CWBDs with specific bacterial host genera in our dataset, together with the literature showing that many of these CWBDs are able to recognize ligands that are specific traits of the related bacterial hosts. For example, *SH3_5* in staphylococcal phages, *CW_binding_1* in *Streptococcus* Mitis group phages, *PSA_CBD* in listeriophages, or *PG−binding_3* in G−. This also manifests the genetic trading between host and parasite, since many of those CWBDs, as well as their bacterial ligands, are also used by the bacterial host surface proteins.
c. The differential appearance of EAD families within phages that infect bacteria with a certain chemotype, which suggests an adaptation of the enzyme to the structure of the specific peptidoglycan it has to degrade. This is notable in the case of peptidases. The somewhat wide range of peptidases identified within our data set is mainly distributed among phages infecting bacteria with subtype A1 peptidoglycan. In phages that infect subtype A3 bacteria, the most common EAD is *CHAP*, which has been shown to function either as NAM-amidase or as endopeptidase, in any case, specific for A3 peptidoglycan.
d. The remarkably differential distribution of domain families among phages that infect either G+ or G−, together with the association of such domains with different physicochemical properties.
e. The differential physicochemical properties between lysins from G+ and G− that, conversely, allows to predict the Gram group of the bacterial host of a given lysin based on its sequence. In this work, the trait of a positively charged patch at a C-terminal position was found to be widespread among lysins from G− bacteria infecting phages. Such trait has been previously related with an improved ability to interact with the G− OM, and might be a ‘substitutive’ of the typical G+ CWBDs. The higher values of other physicochemical variables in G− (aliphatic index, hydrophobic moment) also suggest an analogy of certain structural segments of G− lysins with AMPs.

These observations have clear implications on the design and development of lysin-based antimicrobials, from rational search (or design) of novel lysin parts to deriving AMPs from lysins sequences. A possible setup in which specific bacterial infections are tackled in a personalized manner based on a knowledge-driven, highly efficient synthetic biology platform for lysin-based antimicrobials production can be envisioned in a near future. The conclusions of this work can contribute to the consolidation of such a framework, together with the cutting-edge research currently being carried out in the field.

## METHODS

### Sequence database construction and curation

Phage genomes were retrieved from NCBI nucleotide database by searching phage complete genomes constrained to several bacterial taxa of interest, mainly selected by clinical or epidemiological importance and availability. Those genomes were screened for gene products whose annotations could suggest them to be lytic enzymes. Therefore, keywords such as ‘lysin’, ‘lysozyme’, ‘murein’, ‘amidase’, ‘cell wall hydrolase’, ‘peptidase’ or ‘peptidoglycan’ were used as inclusion criteria, while ‘structural’, ‘tail’, ‘holin’, ‘baseplate’ or ‘virion protein’ were used as exclusion terms to try and avoid misidentifications. Associated information such as taxon of the bacterial host, aa sequence, annotations, phage denomination, and protein/genome unique identifiers were also added into the database.

Curation included: 1) a sequence length cutoff, established with a minimum of 50 and a maximum of 550 aa residues; 2) a sequence identity cutoff using CD-HIT (68) with default parameters and a 98% identity cutoff value to avoid redundant entries; 3) examination with PfamScan (expectation value cutoff = 10) (69, 70) to rule out sequences where no relevant significant hits were found (*i.e.,* where no functional domains that would plausibly appear within phage lysins were detected); 4) bacterial host genus assignation to each entry based on literature and genome annotations). The complete lysins collection and PF hits are available as Table S1 in the supplemental material and at Digital.CSIC (71).

### Physicochemical properties prediction and analysis

Prediction of physicochemical properties (net charge, aliphatic index, hydrophobicity, hydrophobic moment) based on the aa sequences retrieved were performed using the R package ‘Peptides’ implementation (72). Dawson’s pK_a_ scale was used for prediction of net charge assuming pH = 7.0 (73); hydrophobicity scale was that proposed by Kyte and Doolittle (74) and hydrophobic moment was calculated as previously proposed (75) with a specified rotational angle of 100° (recommended angle for α-helix structures). An average value of the hydrophobic moment of each of the possible 11-aa helices within a given sequence is given whenever noted. Such properties were predicted in the whole sequences, in sequences quartiles (contiguous fragments of sequences that account in length each for a quarter of the whole sequence) or in peptides of 11 aa length to provide either a global vision or more local information.

A random forest algorithm was used to check the ability of physicochemical properties to predict lysin sequences as from a G+ or G− infecting phage. R package ‘caret’ was employed for creating, fitting and testing the random forest, and further analyses on the model (ROC curve, MDS plotting) were performed using packages ‘pROC’ and ‘randomForest’. The dataset was randomly partitioned into a training subset (75% of all entries) and a testing subset. The training subset was used to fit the random forest parameters (namely, the randomly selected variables for each node, which was fixed in 4) by a 5-fold cross-validation with 3 repeats. Then the constructed random forest was validated using the previously defined testing subset.

### Sequence similarity networks

SSNs were generated for visually assessing the similarity clustering of sequence sets. For this purpose, the Enzyme Similarity Tool from the Enzyme Function Initiative server (EFI-EST) was employed (76). Briefly, this tool performs a local alignment from which every possible pair of sequences receives a score similar to the E-value obtained from a typical BLAST analysis. A threshold score value was selected for each SSN so that below such threshold sequence pairs were considered nonsimilar and, therefore, the pair would not be connected in the resulting representation. Scores were selected so that sequence pairs whose similarity was below 30-40% were deemed non-similar. The SSN graphs were produced Cytoscape 3 with yFiles organic layout (77).

### Statistical analysis

Default methods for data representation implemented in ‘ggplot2’ R package such as kernel density estimation or GAM smoothing were used throughout this work for data visualization (78). For comparison of non-normal, heteroskedastic data populations, robust statistical methods were used (79). Specifically, a generalization of Welch’s test with trimmed means (default trimming level γ = 0.20) was used with Bonferroni adjustment when multiple comparisons were performed. Effect sizes were estimated according to Wilcox and Tian’s ζ (80). A general interpretation for ζ is given in the previous reference, being values of around 0.10 a small effect size, around 0.30 a medium effect and 0.50 and above a large one. A *p*-value ≤ 0.05 was considered significant. All robust methods were used from the implementation in R Package ‘WRS2’ (81).

### Data Availability

All data used throughout this work are available at Digital.CSIC repository (http://hdl.handle.net/10261/221469) and in Supplemental Material.

## SUPPLEMENTAL MATERIAL

**TABLE S1.** Accession numbers, sequences and PF domains predicted for the lysins data set constructed in this work.

**TABLE S2.** Traceability information and yield of the curation process.

**FIG S1.** Schematic architecture of the LyA autolysin of *S. pneumoniae*.

**TABLE S3.** Information on the PF families found within the lysins sequences database.

**FIG S2.** Heatmap depicting PF hits distribution among different streptococci.

**FIG S3.** Sequence similarity networks (SSNs) of the PF hits in our dataset corresponding to different domain families.

**FIG S4.** Local computation of physicochemical properties in lysins from G− infecting phages classified according to EAD predictions.

## ACKNOWLEDGMENTS

This study was funded by a grant from the Ministerio de Economía y Competitividad (MINECO-FEDER, SAF2017-88664-R). Additional funding was provided by the Centro de Investigación Biomédica en Red de Enfermedades Respiratorias (CIBERES), an initiative of the Instituto de Salud Carlos III. Roberto Vázquez was the recipient of a predoctoral fellowship from CIBERES.

The authors gratefully acknowledge Guillermo Padilla for his fundamental advice in statistical data treatment and representation.

The authors have declared that no competing interests exist.

Authors contribution statement: RV and PG conceptualization; RV performed the analysis and constructed the database; RV and EG data curation; RV wrote the original draft of the paper; all authors read, edited and approved the final manuscript.

